# Understanding Brain Aging Through Behavioural and Microglial Changes: A Lifespan Approach

**DOI:** 10.64898/2026.01.04.697400

**Authors:** María Milagros Sisti, Sofia Cervellini, Facundo Peralta, Franco Juan-Cruz Dolcetti, Osvaldo Martin Basmadjian, Macarena Lorena Herrera, Maria José Bellini

## Abstract

Aging is characterized by progressive physiological decline linked to inflammaging, a chronic low-grade inflammatory state. This study investigates age-related behavioural changes and their correlation with microglial function in female Sprague Dawley rats across their lifespan. Using a longitudinal design at 2, 6, 12, and 24 months of age, we assessed motor performance, mood-related behaviours, and spatial cognition alongside microglial morphometric analysis in key brain regions. Results showed that motor and cognitive performance began to decline significantly at 12 months, with severe impairments and depressive-like behaviours appearing by 24 months. These deficits were paralleled by progressive gliosis in the hippocampus and striatum. Microglial morphometric analysis further indicated a more reactive, region-dependent phenotype, with cells in the striatum adopting a smaller area, reduced perimeter, and fewer intersections.

These findings provide compelling evidence for significant age-dependent deterioration in motor performance, mood regulation, and cognitive abilities in female rats. Our data strongly suggest an underlying progression of neurobiological changes, with microglial dysfunction and neuroinflammation being central candidates. This research contributes valuable insights into the cellular correlates of behavioural decline across the female lifespan, serving as a reference for the multifaceted changes that occur during normal aging.

## 1. Introduction

Aging is a progressive, time-dependent process characterized by loss of physiological function and reproductive capacity, driven by cumulative cellular and tissue changes that destabilize homeostasis and ultimately compromise organismal survival (López-Otín et al., 2023; Moldakozhayev & Gladyshev, 2023). These changes encompass genomic instability, telomere attrition, epigenetic alterations, loss of proteostasis, deregulated nutrient sensing, mitochondrial dysfunction, cellular senescence, stem cell exhaustion, and altered intercellular communication, among other emerging hallmarks (Guo et al., 2022a; López-Otín et al., 2023). A central feature of this landscape is a chronic, low-grade, sterile inflammatory state termed “inflammaging,” which is now recognized as a key driver of age-related morbidity and mortality (Andonian et al., 2025; Ferrucci & Fabbri, 2018). At a central nervous system (CNS) level, aging affects synaptic connectivity, neurotransmitter synthesis, and glial cell function, representing the main cause of age-associated cognitive, motor, and behavioural decline (Blinkouskaya et al., 2021). Within the CNS, inflammaging is tightly linked to functional loss or maladaptive reprogramming of glial cells, particularly microglia and astrocytes, which progressively shift from homeostatic to dysfunctional and often pro-inflammatory states (Hanslik et al., 2021; Salas et al., 2020). Microglia, the resident immune cells of the brain, normally survey the parenchyma, clear debris and misfolded proteins, and secrete cytokines that can adopt either anti-inflammatory or pro-inflammatory profiles depending on context (Salas et al., 2020). With age, microglia undergo characteristic morphological remodelling, show enhanced basal pro-inflammatory signalling, exhibit reduced phagocytic capacity, and develop region-specific dysfunction, ultimately promoting chronic neuroinflammation, impaired learning and memory, and increased vulnerability to neurodegenerative disease (Hanslik et al., 2021; Püntener et al., 2012; Salas et al., 2020).

Across the lifespan, multiple studies have described transcriptomic, numerical, and morphological shifts in both microglia and astrocytes. Our own work has identified region-specific differences in glial cell number, morphology, and transcriptomic profile in senile rats and in a parkinsonism model, with clear implications for both motor and cognitive performance (Falomir-Lockhart et al., 2022; Herrera et al., 2024). Pioneering work in adult mice demonstrated marked heterogeneity in microglial distribution and morphology across brain regions, but this analysis was restricted to a relatively narrow age range (Prinz & Priller, 2014; Tan et al., 2020). Since rats are among the most widely used models in behavioural and neurobiological research and given documented species differences between mice and rats in neuroinflammatory and injury responses (Byrnes et al., 2010; Du et al., n.d.), a systematic characterization of age-related glial changes in naïve rats is warranted. Inspired by the concept of temporal progression, a central goal of this research is to identify an optimal window for therapeutic intervention, especially in females who exhibit extended longevity. We aimed to address this knowledge gap by studying the number, distribution, and morphology of microglia in key brain regions and to relate these parameters to motor and cognitive performance in rats at different ages. A better understanding of these physiological, age-dependent glial trajectories may help to identify early cellular correlates of behavioural decline and inform strategies to prevent or delay neurodegenerative disease.

## 2. Materials and Methods

### 2.1 Animals

Female Sprague Dawley rats, of 2, 6, 12 and 24 months old from our animal facilities were used. Animals were maintained in a temperature-controlled room (22 ± 2°C) on a 12-hour light, 12-hour dark cycle and received food and water ad libitum through all the experiments. Animal care and procedures were in accordance with the Animal Welfare Guideline of NIH (INIBIOLPS’s Animal Welfare Assurance No A5647-01) and were approved by the Universidad Nacional de La Plata Committee on Laboratory Animals (CICUAL; Protocol #P05-03-2017).

### 2.2 Experimental Protocol

This study aimed to evaluate age-related changes in behaviour and microglia using a longitudinal design with independent cohorts of animals at four distinct time points: 2, 6, 12, and 24 months. Independent groups were selected for each time point to avoid repeated testing effects and minimize stress, ensuring that observations reflected true age-related differences rather than cumulative experimental influence. Behavioural assessments were organized according to the valence and severity of the tasks, progressing from low-stress to higher-stress paradigms. Tests were spaced out with adequate rest intervals to reduce fatigue and prioritize animal welfare, in compliance with ethical guidelines for laboratory animal care (Figure 1). This approach allowed for reliable behavioural profiling while safeguarding physiological integrity. At the conclusion of each time point, animals were euthanized by decapitation under deep anaesthesia under deep anaesthesia to preserve tissue quality. Brains were carefully extracted, and coronal sections were prepared for immunohistochemical analysis, enabling a comprehensive evaluation of structural and functional changes across the lifespan.

**Figure 1:**
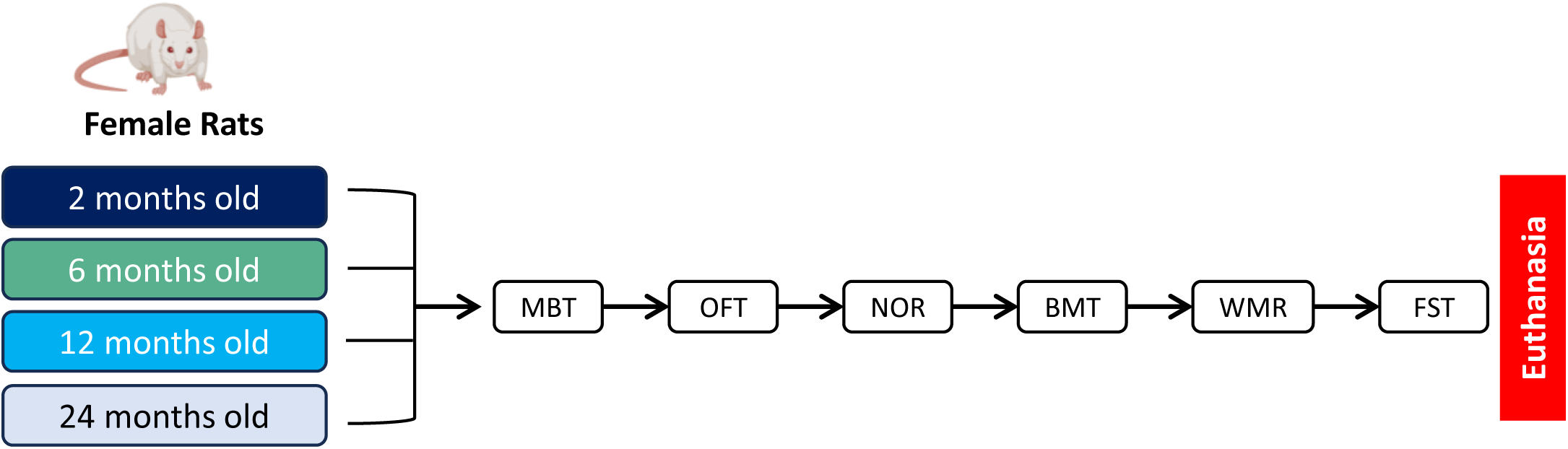
Schematic representation for the experimental design of a lifespan approach of female rats. MBT: Marble Burying Test. OFT: Open Field Test. NOR: Novel Object Recognition Test. BMT: Barnes Maze Test. WMR: Wire Mesh Ramp Test. FST: Forced Swim Test.

### 2.3 Behavioural Assessments

Behavioural testing was conducted between 10:00 a.m. and 4:00 p.m. in a controlled environment with low-intensity lighting. All evaluations were performed by two experimenters to maintain consistency. To minimize stress and facilitate habituation, animals were subjected to handling sessions and placed in the experimental room for 40 minutes prior to each test. Each session was recorded using a video camera positioned above the arena, and behavioural data were analysed offline manually by a blinded observer. All arenas were cleaned with 70% ethanol between trials to eliminate odour cues. For each time point, cohorts were evenly distributed across the four experimental groups and tested simultaneously under identical room conditions.5

#### Wire Mesh Ramp

In this assay, we implemented a modified version of the wire mesh ramp test, originally described by (Nishida et al., 2011), as a measure of grip strength. Each animal completed three trials with a 15s rest between trials, and the median value was used for analysis. The apparatus consisted of a ramp fitted with a central wire-mesh strip (90 cm long × 42 cm wide) positioned at a 70° incline and partially submerged in water to a depth of 15 cm. Animals were placed on the wire-mesh surface, and the time until they lost grip and slid into the water was recorded (Figure 2a). A maximum cutoff time of 120 seconds was set for each trial, and all measurements were performed in triplicate.

**Figure 2:**
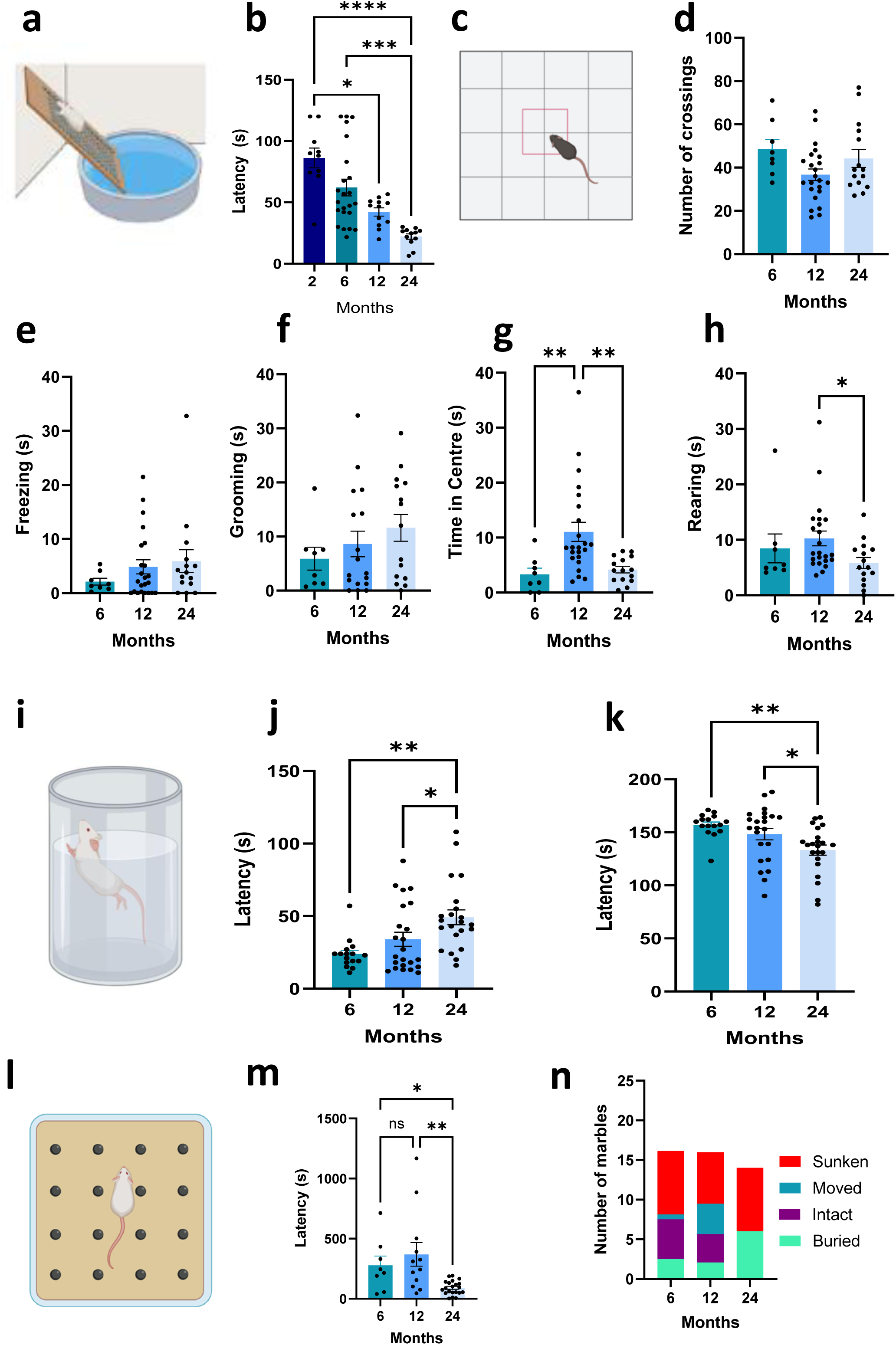
Impact of aging in motor performance and mood-like disorders. (a) Schematic representation of the Wire Mesh Ramp Test (b) Average Latency (seconds) to fall of the WMR test. Kruskal-Wallis test. Dunńs multiple comparison test. N=10 (2 months), N=23 (6 months), N=12 (12 months), N=12 (24 months). (c) Schematic representation of the Open Field Test (d) Open Field: Number of crossings. (e) Open Field: Freezing time (seconds). (f) Open Field: Grooming (seconds). (g) Open Field: Time spent in the centre squares of the arena (seconds). (h) Open Field: Rearing time (seconds). For all the OF measures we performed Kruskal-Wallis test and Dunńs multiple comparison as a *pos hoc* test. N=8 (6 months), N=23 (12 months), N=15 (24 months). (i) Schematic representation of the Forced Swim Test (j) Forced Swim Test: Floting time (seconds) (k) Forced Swim Test: Swimming time (seconds). For all the FST measures we performed Kruskal-Wallis test and Dunńs multiple comparison as a *pos hoc* test. N=16 (6 months), N=23 (12 months), N=22 (24 months). (l) Schematic representation of Marble Burying test. (m) Marble Burying Test: Latency time (seconds) to start digging. Kruskal-Wallis test and Dunńs multiple comparison as a *pos hoc* test. (n) Marble Burying Test: Number of buried, intact, moved and sunken marbles. Two-way ANOVA test. N=8 (6 months), N=12 (12 months), N=20 (24 months). Data are expressed as mean ± SEM. (*P< 0.05; **P=0.01, ***P=0.001; ****P<0.001)

#### Open Field

The Open Field test (OF) is used to determine locomotor activity, exploration habits and anxiety-like behaviour (Voikar et al., 2023). It consists of a square arena (65 cm long x 65 wide x 45 cm depth) whose floor is divided in 16 equal squares (Figure 2c). The animals were placed in the centre of the box and allowed to explore the arena freely for 5 min. We measured i) time spent by the animal grooming or scratching; ii) time spent by the animal exploring in two feet; iii) time the animal is immobile; iv) number of lines crossed with its four legs; and the time spent by the animal in the central square with its four legs.

#### Forced Swim Test

We employed the Forced Swim test (FST), commonly used for the study of depressive-like behaviour in rodents. Active and passive behaviour were evaluated when animals were forced to swim in a cylinder from which they could not escape. A vessel of 30 cm diameter and 40 cm depth was used, with clean water at 26 ± 1°C up to a level of 30 cm (Figure 2i). On day 1 (habituation) the rats were placed in the vessel for 10 min. On day 2 (FST), a 5 min test was performed. Total swimming time (active behaviour) and floating time (depressive-like behaviour) were measured.

#### Marble Burying Test

The Marble Burying (MB) test is commonly used to estimate defensive burial in rodents. It is based on the observation that rodents bury exhibits in their bedding, like glass marbles. Rats were individually placed in a housing cage (30 × 30 × 17 cm) with 5 cm of fresh wood shavings. Sixteen glass marbles (1.5 cm in diameter, arranged in a 4 × 4 pattern) were evenly spaced over the surface (Figure 2l). The number of intact, moved, covered, or buried marbles was analysed during a 30-min period. A marble was considered covered if at least 1/3 of it was covered with shavings and buried if it was completely covered by shavings and at not less than 3 cm from the floor of the cage. In addition, the time taken to start the first dig or bury -latency time (seconds)- was evaluated.

#### Novel Object Recognition

The Novel Object Recognition (NOR) test assesses short- and long-term memory based on an animal’s ability to recognize and explore a novel object. In this study we used a protocol of four stages: i) Habituation, one day before training, animals are placed for 5 min in a square acrylic box (65 × 65 × 45 cm) with a black floor; ii) Training, where animals explore two identical objects (A); iii) Short-Term Memory (NOR 2), where after 90 min, one object is replaced with a novel object (B) and iv) Long-Term Memory (NOR 3), where after 24 h, one object is replaced with a different novel object (C). In all the sessions, female rats explored for 5 min and the exploration time for each object was recorded (Figure 3a).

**Figure 3:**
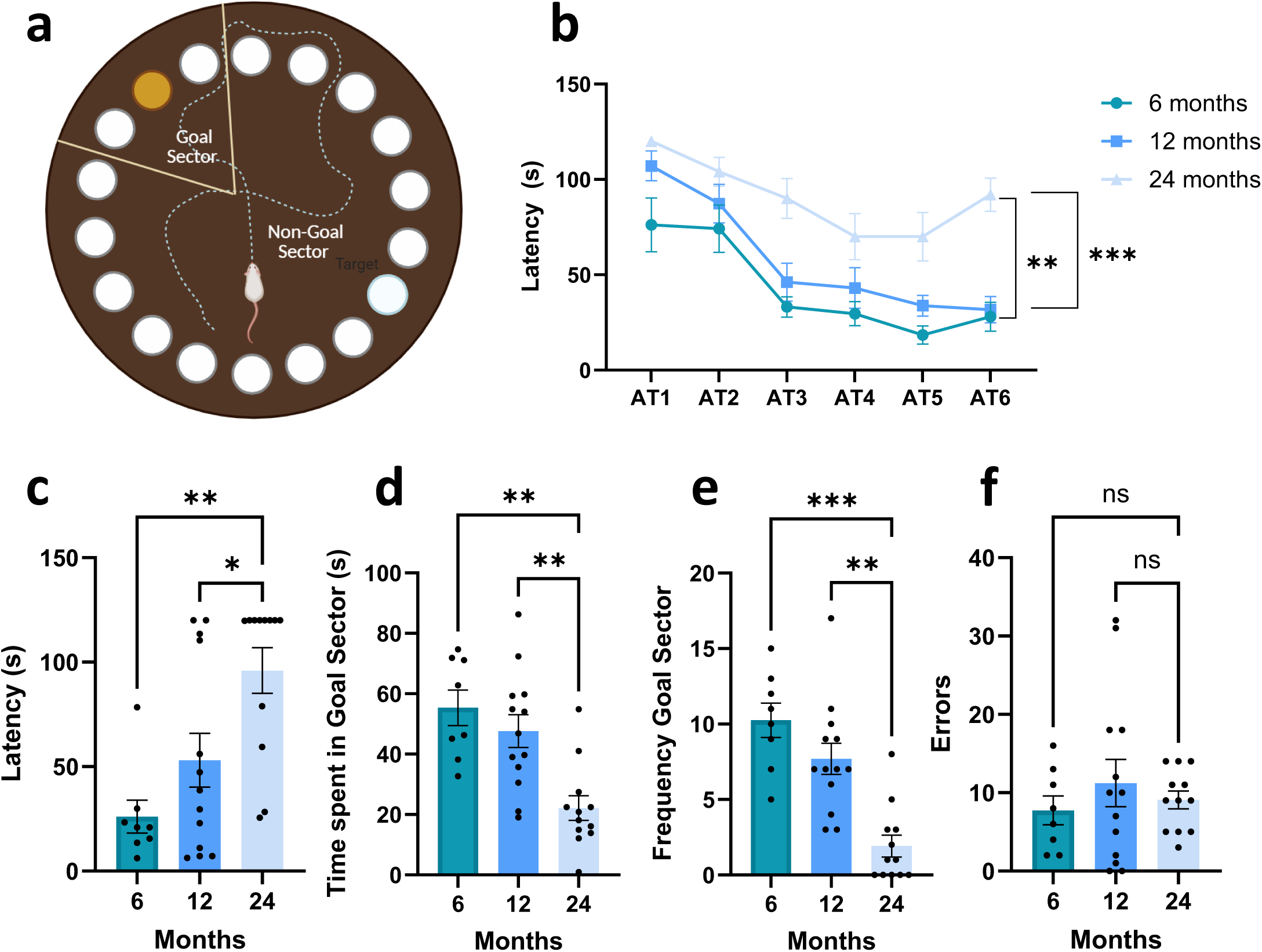
Impact of aging in cognitive performance. (a) Schematic representation of the Barnes Maze Test (b) Latency time (seconds) during the acquisition trials (AT). Two-way ANOVA test and Tukeýs multiple comparison test. N=8 (6 months), N=13 (12 months), N=12 (24 months) (c) Probe trial: Latency (seconds) to reach the goal sector. (d) Probe Trial: Time spent in the goal sector (seconds) (e) Probe Trial: Number of visits to the goal sectos (frequency) (f) Probe Trial: Number of errors For all the PT measures we performed Kruskal-Wallis test and Dunńs multiple comparison as a *pos hoc* test. N=8 (6 months), N=13 (12 months), N=12 (24 months). Data are expressed as mean ± SEM. (*P< 0.05; **P=0.01, ***P=0.001; ****P<0.001)

#### Barnes Maze Test

For the Barnes Maze (BM) test, we followed the protocol by Morel et al., (Morel et al., 2015) to assess spatial memory using a raised black acrylic circular platform (122 cm diameter, 108 cm high) with 20 evenly spaced holes (10 cm diameter). Only one hole connects to a removable black escape box (38.7 × 12.1 × 14.2 cm). Rats start in an opaque white cylinder (25 cm diameter, 20 cm height) placed at the centre. Four distal visual cues are positioned 50 cm from the platform. Hole 0 (escape hole) remains fixed; others are numbered clockwise (1–10) and counterclockwise (−1 to -9). A 90 dB white noise and a 500 W light serve as aversive stimuli (Figure 3XX). We performed the sessions of habituation where rats were placed in the start cylinder and escape box for 120 s each, training or Acquisition Trials (AT) where rats started in the cylinder for 30 s; then aversive stimuli are activated, and rats explore for 120 s to find the escape hole and the Probe Trial (PT): Same as AT, but escape box was removed; rats explored for 120 s to assess memory retention. We measured the following variables: i) Latency: Time to enter the escape box (AT) or first exploration of escape hole (PT); ii) Goal Sector Exploration (GS): Number of explorations of holes -1, 0, and 1; iii) Time in Goal Quadrant: Time spent in the quadrant containing the escape hole (PT); iv) Non-Goal Hole Exploration: Explorations of holes other than the escape hole (PT).

#### Sample Processing

Rats were anesthetized with isoflurane (Baxter, USA) and sacrificed by decapitation. Brains were removed and split into hemispheres: one for molecular analysis and the other for immunohistochemistry (IHC). Hemispheres for IHC were post-fixed in 4% paraformaldehyde (pH 7.4) overnight at 4°C, then stored in cryoprotective solution (30% ethylene glycol, 30% sucrose in 0.1M PB, pH 7.4) at −20°C. Coronal sections (40μm) were cut using a Leica vibratome and organized into eight series, each containing non-contiguous sections spanning different brain regions.

#### Immunohistochemistry

Sections were washed in 0.1 M phosphate buffer (PB, pH 7.4) with 0.3% Triton X-100. Endogenous peroxidase was quenched for 10 min in 3% H_2_O_2_/50% methanol. Non-specific binding was blocked with PB + 0.3% Triton X-100 + 5% serum (species-specific). After washes, sections were incubated overnight at 4°C with Iba1 rabbit polyclonal antibody (1:1500; Wako, Ref. 016-20001). Next, they were incubated for 2h at room temperature with biotinylated anti-rabbit secondary antibody (1:1000; Vector BA-1000), followed by 90min in ABC complex (1:500; Vector PK-6100). Immunoreactivity was visualized using DAB and 0.01% H_2_O_2_ in PB. Sections were mounted on gelatine-coated slides, dehydrated, and coverslipped with Canada balsam for image analysis.

#### Morphometric Analysis

The number of Iba1 immunopositive cells in the hippocampus (dentate gyrus and *cornus ammonis* CA1) and *striatum* were assessed. Areas of interest were defined in accordance with the rat brain atlas of Paxinos and Watson^7^. Imaging of sections was carried out using an Olympus BX-51 direct microscope connected to an Olympus DP70 CCD video camera. Images of all focal planes were captured at 60X, analyzing a total of 10 fields for hippocampal areas and 30 fields for the striatum per rat. Iba1 immunopositive cells were quantified automatically, using a macro from a previous study Herrera et al., (Herrera et al., 2024), and running on the FIJI-ImageJ software.

#### Microglia Reactivity Analysis

Microglia images were processed in FIJI-ImageJ software using two steps: segmentation and analysis. i) <u>Segmentation</u>: Grayscale images were converted to binary using Li and percentile algorithms. Filters (unsharp, band-pass) enhanced contrast and removed noise. Skeletonization and boundary masks prevented particle merging, producing clean binary masks for analysis. ii) <u>Analysis for microglia</u>: number of reactive cells, average area occupied per cell, perimeter, round, circularity, solidity, and Sholl analysis (ending radius, number of total intersections per cell and intersections per 5 μm up to 100 μm).

#### Statistical Analysis

The statistical analysis was conducted using GraphPad Prism software version 10.0.3 for Windows (GraphPad Software, Inc, San Diego, CA, USA). To assess the normality of the data we used the Kolmogorov-Smirnov and Shapiro-Wilk tests and to assess homoscedasticity we used the Levene and Brown-Forsythe tests. Group comparisons were performed using one-way analysis of variance (ANOVA). Upon observing statistical significance, Dunn’s or Tukey’s post hoc test was employed. Significance was considered at P < 0.05. Specific tests and significance for each case are detailed in the figure captions.

## 3. Results

### 3.1 Age-related impact in motor performance and mood-like disorders in female rats

In Figure 2a-b, graphs show the strength assessment for female rats through different stages. Female rats exhibited a gradual decrease in grip strength (H _(3)_ =30.52; P<0.0001, Figure 2b). This difference was statistically significant at 12 months (P=0.034) and exacerbated at 24 months compared with 2 months (P<0.0001) and 6 months (P=0.001). Furthermore, regarding the exploration of the open field, we could not observe significant difference in the number of lines crossed (H_(2)_=4.86; P=0.08, Figure 2d), the freezing time (H_(2)_=4.86; P=0.08, Figure 2e), nor the grooming time (H_(2)_=1.40; P=0.49, Figure 2f). However, the time spent in the centre peaks at 12 months (H _(2)_ =15.92; P=0.003, 2g), both younger (6 months) (P=0.0031) and older (24 months) (P=0.0040) groups spend significantly less time in the centre than the 12-month group. Additionally, animals of 24 months spent less time doing rearing (H _(2)_ =6.18; P=0.045, 2h). All this suggests an age-dependent change in exploration behaviour, starting at mid-age. On the other part, to assess depressive-like symptoms we performed the force swimming test, where the three groups had an increment of floating times with a peak at 24 months (H_(2)_=13.23; P=0.0013, 2j), with statistical differences with 6 (P=0.0020) and 12 months (P=0203), respectively; and conversely the lowest swimming times (H_(2)_=11.90; P=0.0026, 2k) with also significant differences at 6 and 12 months (P=0.0033 and P=0.0375) respectively. Finally, in the marble burying test, we observed a significative decreased in the latency to start digging (H_(2)_=12.59; P=0.0018, 2m) in the group of 24 months compared to the 6 months (P=0.04) and 12 months (P=0.0036), and with the most number of buried and sunken marbles (F_(6, 148)_=5.858; P<0.0001, 2n).

### 3.2 Age-related impact in cognitive performance in female rats

Performance in the Barnes maze differed across age groups, indicating age-dependent changes in spatial learning and memory (Figure 3). Analysis of learning curves across acquisition training days (F_(5, 180)_ =20.13; P<0.0001, 3b) revealed that 6-month-old female exhibited a robust improvement over time, with a progressive decrease in latency across trials. In contrast, 12-month-old animals showed a slower rate of improvement, while 24-month-old animals demonstrated minimal learning compared to 6- and 12-months groups (P=0.0013 and P=0.0003), with performance remaining relatively stable or only modestly improved across days. Notably, age differences in latency times became more pronounced during later training sessions, indicating reduced learning efficiency in older animals (F _(2, 180)_ =39.28; P<0.0001, 3b). In addition, during the ATs, there a significant interaction of the age differences over time in the variable of time spent in the goal sector (F_(10, 180)_=1.98; P=0.0375, S1a) whereas no significant differences were observed in the frequency of visits to the goal sector (F_(10, 180)_=1.467; P=0.155,S1b) nor in the number of errors (F_(10, 180)_=1.67; P=0.090,S1c). During the probe trial, younger animals maintained superior performance, whereas 24-month group showed increased variability and poorer recall, as reflected by a higher latency to find the area of the scape hole (H_(2)_=12.50; P=0.0019,3c), a reduced time spent in the goal sector (H_(2)_=14.18; P=0.0008, 3d) and in the frequency of visits (H_(2)_=12.59; P=0.0018, 3e) and error rates (H_(2)_=18.91; P<0.0001,3f). Overall, these findings suggest a clear age-dependent decline in the BMT, with significant impairments in spatial learning and memory emerging by 24 months of age and intermediate deficits evident at 12 months. Moreover, we wanted to reinforce these findings by assessing short- and long-term memory under a different behavioural paradigm. For this purpose, we used the NOR test (Figure S1d) where during the training session (F _(2,38)_ =1.339; P=0.274, S1e), the only age group that exhibited an object preference was the 6-month group (P=0.023). Moreover, we could not detect any deficits neither the short-term memory (F _(2,38)_ =2.94; P=0.06, S1f) nor the long-term memory (F_(2,38)_ =0.724; P=0.4911, S1g). Taken together, these data suggest that, unlike spatial memory deficits observed in the BMT, object recognition memory is relatively resistant and remaining largely preserved across aging.

### 3.3 Age-related impact in the microglia reactivity of the striatum

To link our behavioural findings in motor performance to underlying neurobiological changes, we performed a stereological analysis of Iba1⁺ cells in the striatum, a key brain region involved in motor control. To assess age-related changes in striatal microglia, we performed immunohistochemical labelling of Iba1 and quantified microglial morphology using stereological and skeleton-based analyses (Figure 4). Representative images (Figure 4a) from young animals revealed microglia with highly ramified processes and extensive territorial coverage, consistent with a homeostatic phenotype. In contrast, aged animals displayed microglia with shorter, thicker, and less complex processes, a morphology commonly associated with microglial senescence and functional decline. Quantitative analysis of striatum slides confirmed an exponential age-dependent increase in the number of microglia cells (H _(3)_ = 9.17; P=0.009, Figure 4b), with statistical significances in the 24-month group compared to 2 and 6 months old (P= 0.0229, P=0.0286) female rats, respectively. These findings in the aged group correlates with a decrease in the average area occupied by cell (H_(3)_=6.43; P=0.069, Figure 4c) compared to the young group of 6 months old (P=0.0286) and a decrease in the perimeter variable (H_(3)_=6.35; P=0.0731, Figure 4d) compared to the young group of 6 months old (P=0.0286). No significant differences were found in the measures of circularity (H _(3)_=5.192; P=0.153, Figure S2a), round (H _(3)_ =2.247; P=0.569, Figure S2b) and solidity (H _(3)_=2.473; P=0.529, Figure S2c). Additionally, Sholl analysis revealed similar final radius of the microglia ramifications (H_(3)_=4.747; P=0.199, Figure 4e), a significant reduction in the total number of intersections in the 24 month group (H_(3)_=851; P=0.011, Figure 4f) compared to the 6 months old group (P=0.0286) and no differences in the number of intersections when we analysed in the range of every 5-µm (F_(57,190)_=0.720; P=0.926, Figure 4g). Overall, the observed age-dependent increase in microglial number and concomitant loss of morphological complexity in the striatum may underlie, at least in part, the motor performance deficits observed in aged animals.

**Figure 4:**
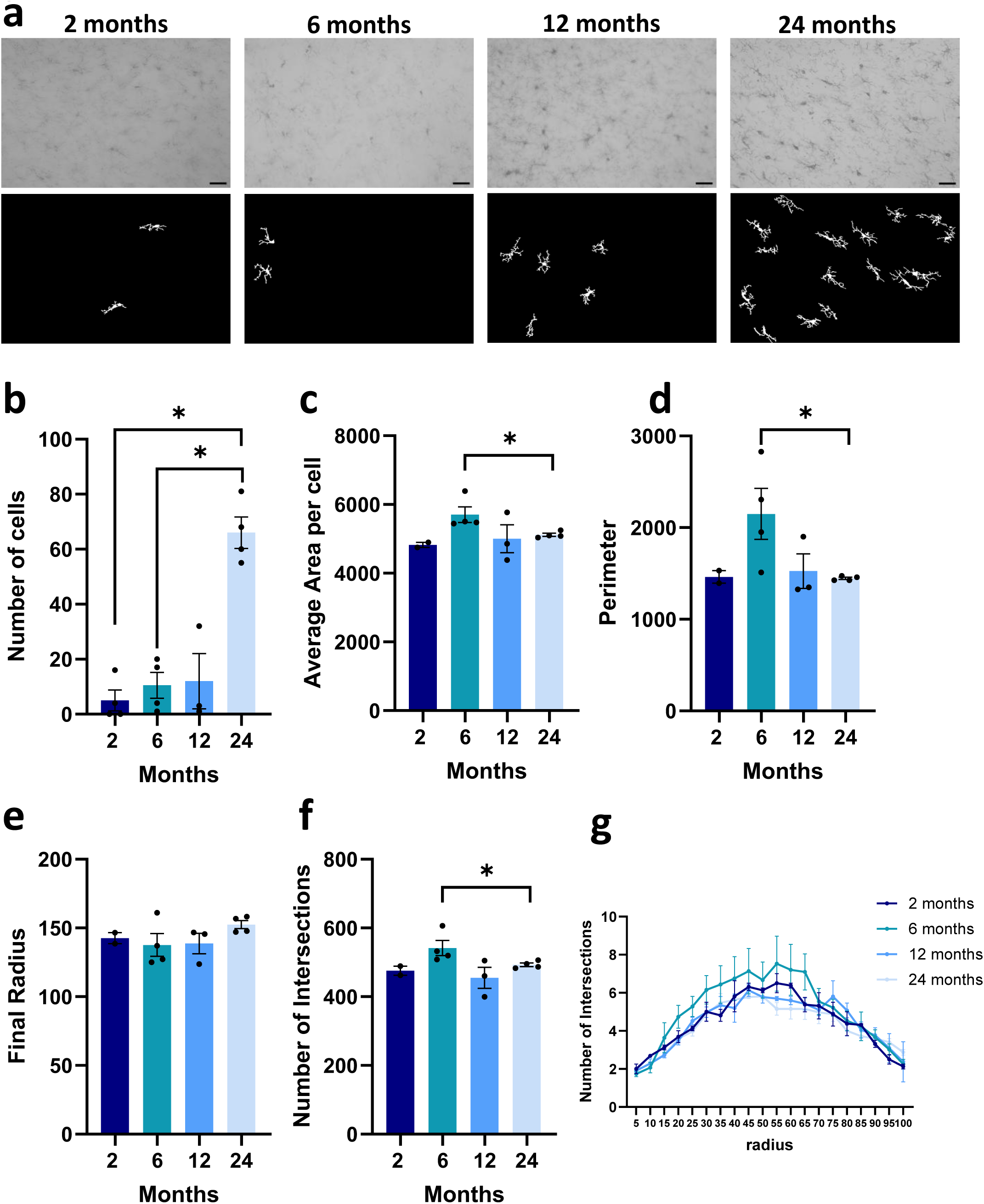
Effect of aging on microglia cells of striatum (ST) (a) Representative original and binary images of microglia cells in each condition at 630X magnification. Bar scale at 500 microns (b) Number of cells: Iba1 immunoreactive cells. (c) Average area occupied per cell: area of selection in square pixels. (d) Perimeter: The length of the outside boundary of the selection. (e) Final radius of intersections (f) Total number of intersections (g) Number of intersections by consecutive radio of 5 microns (fixed at 100 microns). One-way ANOVA test and Tukey as post hoc. Two-way ANOVA of RM. N= 2 (2 months); N=4 (6 months); N=3 (12 months); N=4 (24 months). Data are expressed as mean ± SEM. (*P< 0.05; **P < 0.01; ***P < 0.001).

### 3.4 Age-related impact in the microglia reactivity of the hippocampus

Given the vulnerability of the hippocampus to aging, we examined whether microglial organization within the dentate gyrus is altered across the lifespan. Iba1 immunolabeling combined with stereological and skeleton-based analyses was used to characterize age-related changes in microglial abundance and morphology (Figure 5a). Quantitative assessment revealed a pronounced, age-dependent increase in microglial number within the dentate gyrus (H _(3)_=10.55; P=0.0144, Figure 5b) driven by a significant elevation in 24-month-old females relative to 2-, 6- and 12-month-old groups (P=0.0151, P=0.0247 and P=0.0449, respectively). In contrast to striatum analysis, shape descriptors, including average area per cell (Figure 5c), perimeter (Figure 5d), circularity (Figure S2d), roundness (Figure S2e), and solidity (Figure S2f), were not significantly affected by age. Analysis of process organization using Sholl metrics showed comparable maximal ramification distances across groups (H_(3)_=0.1864; P=0.9798, Figure 5e), similar total number of intersections (H_(3)_=0.2013; P=0.977, Figure 5f) and the spatial distribution of intersections at 5-µm intervals remained unchanged (F_(57,380)_=1.310; P=0.0752, Figure 5f). Together, these results demonstrate that aging in the dentate gyrus is characterized by microgliosis consistent with altered immune surveillance in a region essential for learning and memory. However, when we decided to determine whether aging differentially impacts microglial organization within hippocampal subfields, and examined Iba1+ microglia in the CA1 region, a circuit essential for synaptic integration and memory consolidation (Figure 6a) we could not observed significant differences in all the particle analysis (Figures 6b-d and Figures S2g-i) and in the sholl analysis (Figures 6e-g). Collectively, these results indicate that microglial density and morphological features within the CA1 subregion remain stable across aging.

**Figure 5:**
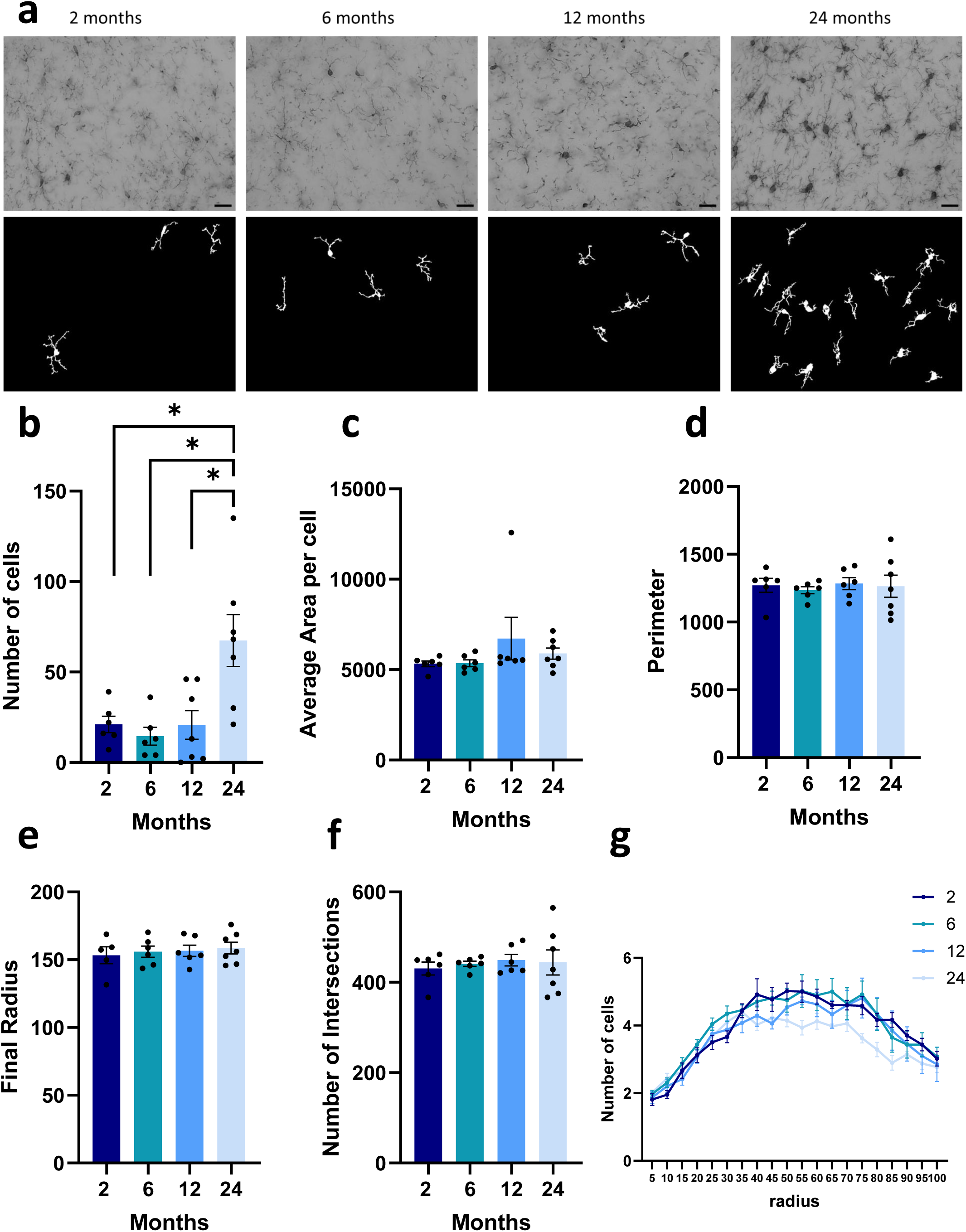
Effect of aging on microglia cells of hippocampal subregion gyrus dentate (GD). (a) Representative original and binary images of microglia cells in each condition at 630X magnification. Bar scale at 500 um (b) Number of cells: Iba1 immunoreactive cells. (c) Average area occupied per cell: area of selection in square pixels. (d) Perimeter: The length of the outside boundary of the selection. (e) Final radius of intersections (f) Total number of intersections (g) Number of intersections by consecutive radio of 5 microns (fixed at 100 microns). One-way ANOVA test and Tukey as post hoc. Two-way ANOVA of RM. N=6 (2 months); N=6 (6 months); N=6 (6 months); N=6 (12 months); N=7 (24 months). Data are expressed as mean ± SEM. (*P< 0.05; **P < 0.01; ***P < 0.001).

**Figure 6:**
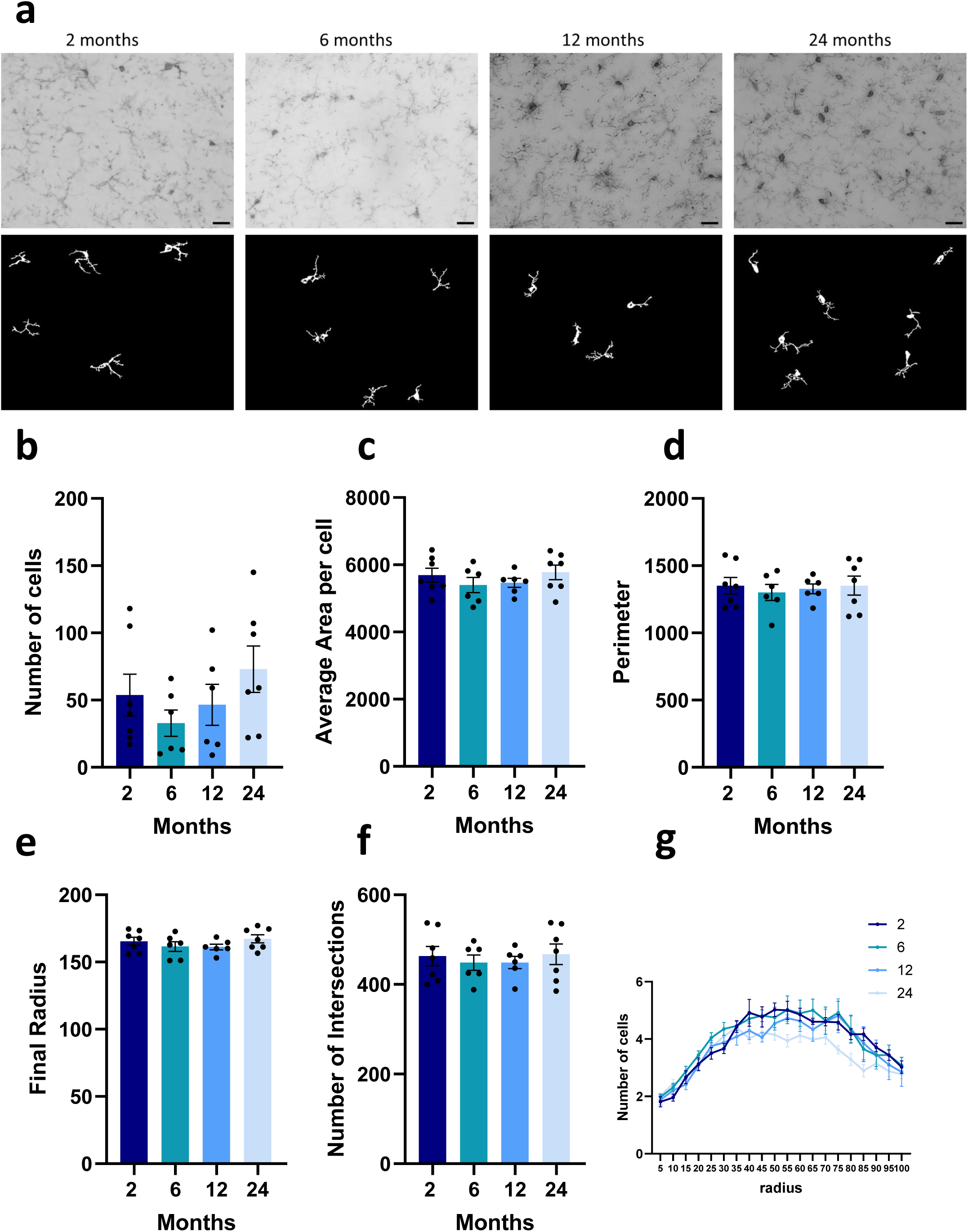
Effect of aging on microglia cells of hippocampal subregion cornice amonis (CA1). (a) Representative original and binary images of microglia cells in each condition at 630X magnification. Bar scale at 500 um (b) Number of cells: Iba1 immunoreactive cells. (c) Average area occupied per cell: area of selection in square pixels. (d) Perimeter: The length of the outside boundary of the selection. (e) Final radius of intersections (f) Total number of intersections (g) Number of intersections by consecutive radio of 5 microns (fixed at 100 microns). One-way ANOVA test and Tukey as post hoc. Two-way ANOVA of RM. N=7 (2 months); N=6 (6 months); N=6 (6 months); N=6 (12 months); N=7 (24 months),. Data are expressed as mean ± SEM. (*P< 0.05; **P < 0.01; ***P < 0.001).

## 4. Discussion

This study employed a lifespan approach to investigate age-related changes in behavioral performance and their potential basis in microglial function in female Sprague Dawley rats. Our findings reveal significant age-dependent declines in motor performance, the emergence of mood-like disorders, and impairments in spatial cognition, particularly evident in older animals. These behavioral alterations, observed across cohorts equivalent to puberal (2 months), young adult (6 months), middle-aged (12 months), and old (24 months) humans (Sengupta, 2013), align with the progressive nature of aging and provide a crucial framework for understanding the physiological and cellular trajectories that contribute to age-associated decline, as proposed in our introduction.

One of our findings from the behavioural assessment is a pronounced decrease in grip strength with advancing age, which becomes statistically significant by 12 months and is further exacerbated at 24 months. This motor deficit is accompanied by reduced rearing behaviour in 24-month-old rats, suggesting a broader impact of aging on motor activity and exploration. While the Open Field test did not show significant differences in locomotion or freezing time, the altered time spent in the central area, peaking at 12 months, appears to follow a hormetic pattern, with 12-month-old animals exhibiting less anxiety as an adaptive response to aging that could not be compensated at 24 months, as described in the literature. (Rattan, 2008; Santoro et al., 2020). These findings hint at age-dependent shifts in exploratory strategies or anxiety-like responses that warrant further investigation. Furthermore, our data indicate an age-related increase in depressive-like behaviours. The Forced Swim Test revealed significantly increased floating times and concurrently decreased swimming times in 24-month-old rats compared to younger cohorts, a classic indicator of behavioural despair. This is further supported by observations in the Marble Burying Test, where older animals showed a decreased latency to start digging and buried/sank more marbles. These combined results strongly suggest an age-dependent predisposition to mood disorders, which are commonly observed in aged human populations (Alexopoulos, 2005, 2019; Herrera et al., 2020).

Cognitive function also displayed a clear age-related decline. Performance in the Barnes Maze, demonstrated a robust improvement in 6-month-old rats during acquisition, which progressively diminished in 12-month-old animals and was severely impaired in 24-month-old rats. The age differences in latency times became more pronounced in later training sessions, underscoring a reduced learning efficiency and memory retention in older cohorts. These findings corroborate the well-established age-associated cognitive decline and align with age-related synaptic remodelling signatures in the rat hippocampus and parahippocampal regions that underlie diminished cognitive reserve (Buss et al., 2021; Horovitz et al., 2025; Mota et al., 2019). Crucially, these observed behavioral deficits are hypothesized to be closely linked to age-dependent alterations in glial cell function, particularly microglia, as posited in our introduction. The progressive declines in motor performance, mood, and cognition are consistent with the concept of “inflammaging” within the central nervous system (Andonian et al., 2025; Ferrucci & Fabbri, 2018; Franceschi et al., 2000, 2007), in which microglia gradually shift from a homeostatic state to a dysfunctional and pro-inflammatory phenotype (Hefendehl et al., 2014; Mosher & Wyss-Coray, 2014; von Bernhardi et al., 2015). Our morphometric analysis of microglia shows progressive gliosis with age, supported by an increased number of microglia in the striatum and both hippocampal areas studied (DG and CA1). Microglia also adopt a more reactive, region-dependent phenotype, as demonstrated by a smaller area, reduced perimeter, and fewer intersections in the striatum but not in the hippocampus. These direct cellular findings support our behavioral observations, providing a more comprehensive understanding of how microglial transformations correlate with specific functional losses across the lifespan. This microglial reprogramming, driven by aging-related molecular mechanisms such as genomic instability, loss of proteostasis, and cellular senescence (Guo et al., 2022b), is a known driver of chronic neuroinflammation, impaired synaptic function, and increased vulnerability to neurodegenerative processes (Andronie-Cioara et al., 2023; Calabrese et al., 2018; Di Benedetto et al., 2017; Mottahedin et al., 2017), which could directly underlie the behavioural changes we report. Moreover, the stepwise behavioural impairments parallel the progression of neuropathological stages that correlate with cognitive decline, as described in Braak staging for Parkinson’s or Alzheimeŕs disease (Braak et al., 2005; Gitzlaff et al., 2024).

The longitudinal design with independent cohorts at distinct time points effectively captured the dynamic nature of aging, minimizing confounding factors such as repeated testing effects. Our specific focus on female rats is particularly relevant given their extended longevity and distinct neuroinflammatory responses compared to males (Austad & Fischer, 2016; Gilmer et al., 2023; Ince et al., 2023; Lemaître et al., 2020), contributing valuable data to a field often dominated by male animal studies (Knufinke et al., 2023; Mauvais-Jarvis et al., 2017). This approach allows for the identification of critical windows where age-related changes become significant, which is essential for developing timely therapeutic interventions.

In conclusion, our study provides compelling evidence of significant age-dependent deterioration in motor performance, mood regulation, and cognitive abilities in female rats. These behavioural trajectories, especially the pronounced declines observed from mid-age onward, strongly suggest an underlying progression of neurobiological changes, with microglial dysfunction and neuroinflammation as central candidates, like molecular hallmarks of aging. Future analyses integrating molecular data will further elucidate the mechanistic correlates of these behavioural deficits, offering insights into potential targets for preventing or delaying age-related neurodegenerative diseases. Taken together, our findings establish a systematic staging framework that serves as a reference for the multifaceted changes occurring during physiological aging.

## Acknowledgements

We would like to thank our technical staff Juan Manuel Lofeudo and Araceli Bigres for their invaluable assistance with animal care.

Schematics figures were created using BioRender software.

We used artificial intelligence to assist with English language improvement and grammatical correction during the drafting of this paper.

## Funding

This research was supported by the UNLP grant 11M219 (MJB).

## Contributions

SC: conceptualization, methodology, investigation, analysis, original draft preparation. MS: visualization, investigation, methodology, analysis. FP: methodology, investigation. FJCD: methodology, investigation. OMB: methodology, analysis. MLH: conceptualization, analysis, data curation, draft preparation, review and editing; MJB: conceptualization, analysis, review and editing, funding, supervision.

## Ethics declarations

### Competing interests

The authors declare no competing interests.

### Data availability

All data associated with this study are present in the paper or the Supplementary Materials. Any additional information that supports the findings reported in this study is available by contacting the corresponding authors upon reasonable request.

---

**Supplementary Figure 1:**
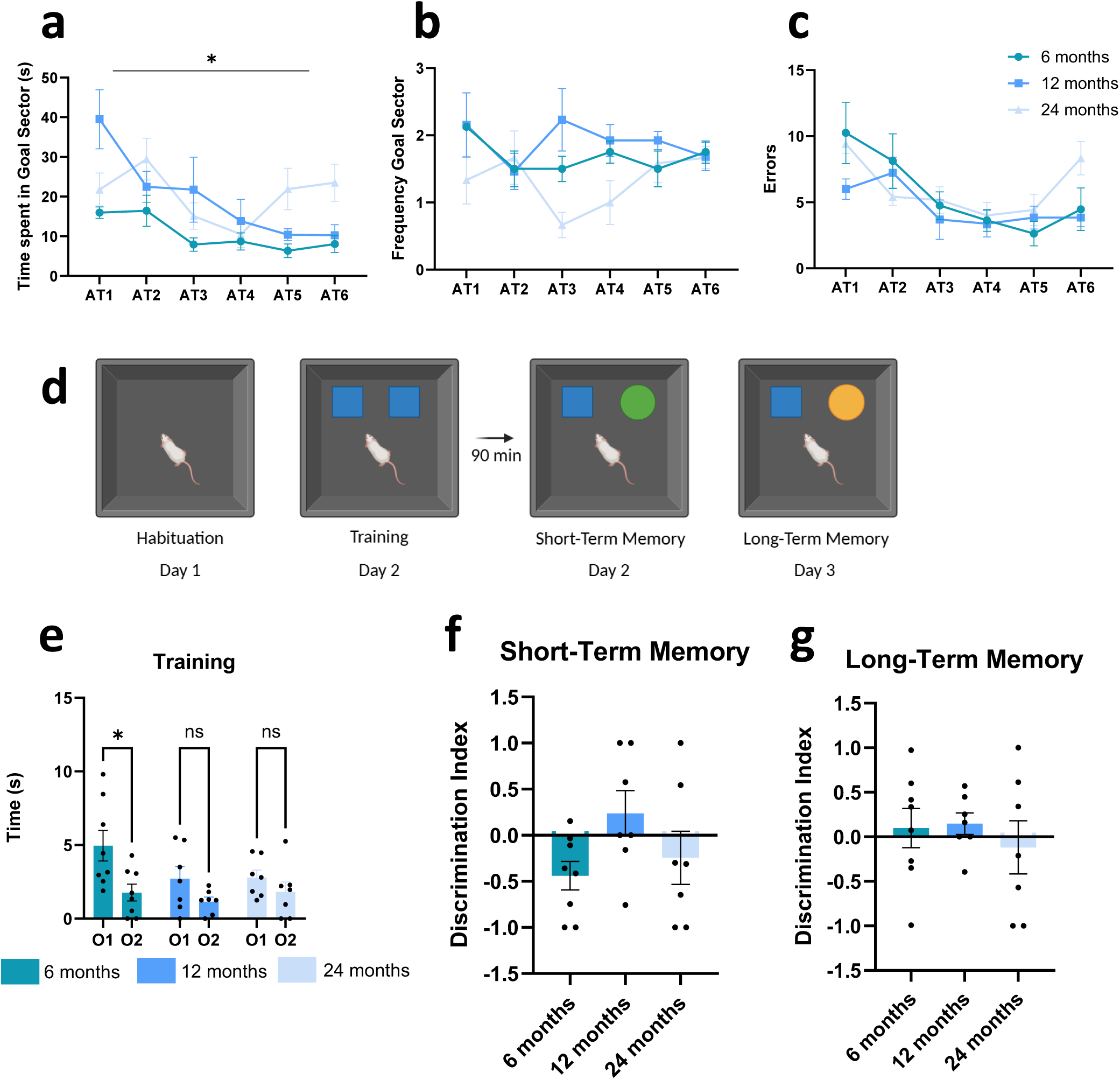
Impact of aging in cognitive performance associated to learning, short- and long-term memory. (a) BMT: Time spent in the goal sector during the acquisition trials. (b) BMT: Number of visits (frequecency) to the goal sector during the acquisition trials. (c) BMT: Number of errors during the acquisition trials. Two-way ANOVA test and Tukeýs multiple comparison test. N=8 (6 months), N=13 (12 months), N=12 (24 months) (d) Schematic representation of the Novel Object Recogntion Test (e) NOR: Training. Two-way ANOVA test and Tukeýs multiple comparison test. Two-way ANOVA test and Tukeýs multiple comparison test. N=8 (6 months), N=7 (12 months), N=7 (24 months) (f) NOR: Discrimination Index to evaluate short-term memory. One-way ANOVA test. (g) NOR: Discrimination Index to evaluate long-term memory. One-way ANOVA test.

**Supplementary Figure 2:**
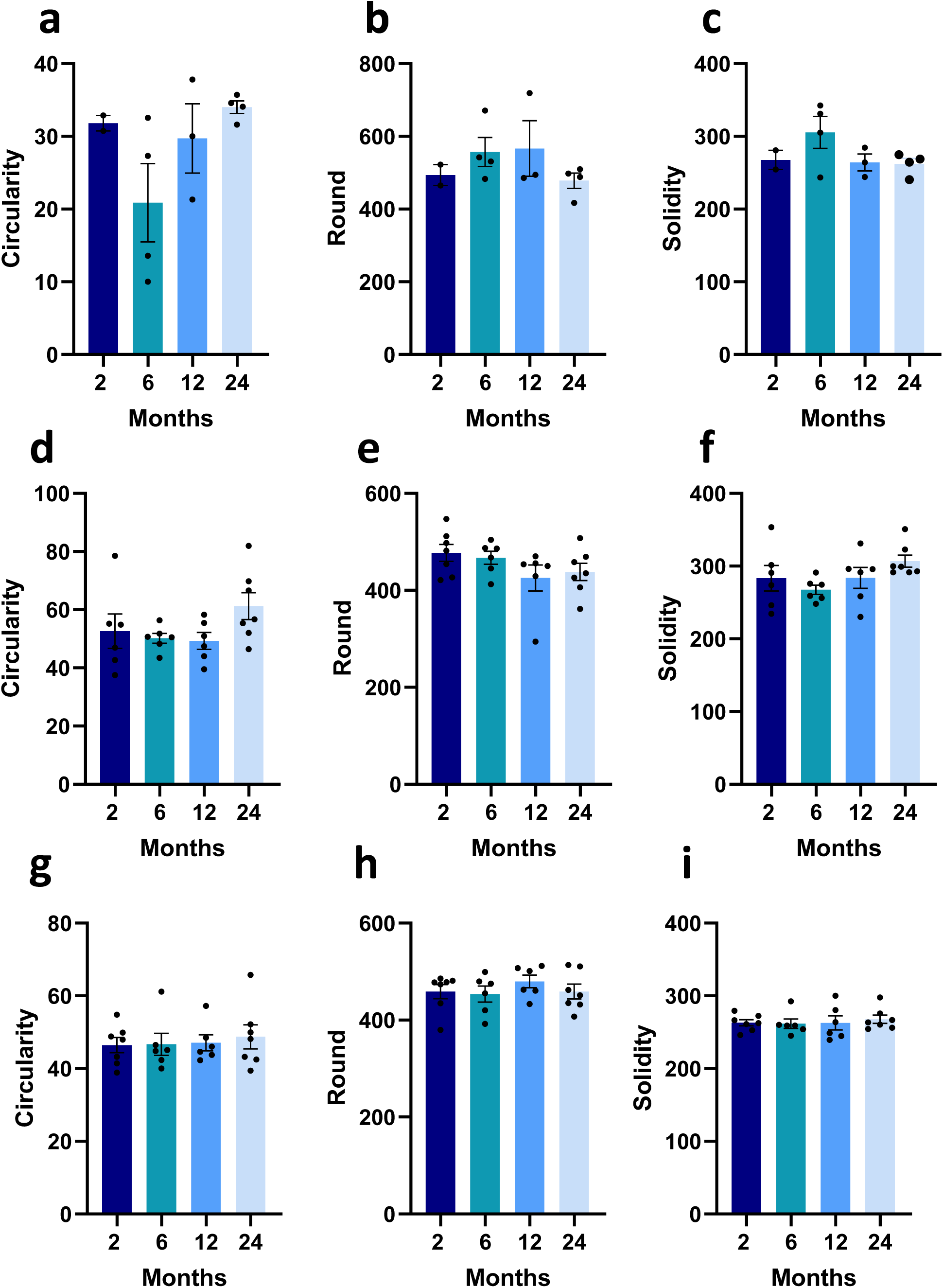
Particle analysis of different brain regions. Striatum. (a) Circularity (b) Roundness: 4*area/(π*major_axis^2), or the inverse of the aspect ratio (c) Solidity One-way ANOVA test and Tukey as post hoc. N= 2 (2 months); N=4 (6 months); N=3 (12 months); N=4 (24 months). Data are expressed as mean ± SEM. Hippocampus: gyrus dentatus (d) Circularity (e) Roundness (f) Solidity One-way ANOVA test and Tukey as post hoc. N=6 (2 months); N=6 (6 months); N=6 (6 months); N=6 (12 months); N=7 (24 months). Data are expressed as mean ± SEM. Hippocampus: CA1 (g) Circularity (h) Roundness (i) Solidity One-way ANOVA test and Tukey as post hoc. N=7 (2 months); N=6 (6 months); N=6 (6 months); N=6 (12 months); N=7 (24 months). Data are expressed as mean ± SEM.

## Notes

### Competing Interest Statement

The authors have declared no competing interest.

## References

1. Alexopoulos, G. S. (2005). Depression in the elderly. The Lancet, 365(9475), 1961–1970. 10.1016/S0140-6736(05)66665-2

2. Alexopoulos, G. S. (2019). Mechanisms and treatment of late-life depression. Translational Psychiatry, 9(1). 10.1038/S41398-019-0514-6

3. Andronie-Cioara, F. L., Ardelean, A. I., Nistor-Cseppento, C. D., Jurcau, A., Jurcau, M. C., Pascalau, N., & Marcu, F. (2023). Molecular Mechanisms of Neuroinflammation in Aging and Alzheimer’s Disease Progression. International Journal of Molecular Sciences, 24(3). 10.3390/IJMS24031869

4. Austad, S. N., & Fischer, K. E. (2016). Sex Differences in Lifespan. Cell Metabolism, 23(6), 1022–1033. 10.1016/J.CMET.2016.05.019

5. Braak, H., Rüb, U., Jansen Steur, E. N. H., Del Tredici, K., & De Vos, R. A. I. (2005). Cognitive status correlates with neuropathologic stage in Parkinson disease. Neurology, 64(8), 1404–1410. 10.1212/01.WNL.0000158422.41380.82;SUBPAGE:STRING:ABSTRACT;R EQUESTEDJOURNAL:JOURNAL:WNL;JOURNAL:JOURNAL:WNL;WGROUP:STRING:PUBLICATION

6. Buss, E. W., Corbett, N. J., Roberts, J. G., Ybarra, N., Musial, T. F., Simkin, D., Molina-Campos, E., Oh, K. J., Nielsen, L. L., Ayala, G. D., Mullen, S. A., Farooqi, A. K., D’Souza, G. X., Hill, C. L., Bean, L. A., Rogalsky, A. E., Russo, M. L., Curlik, D. M., Antion, M. D., … Nicholson, D. A. (2021). Cognitive aging is associated with redistribution of synaptic weights in the hippocampus. Proceedings of the National Academy of Sciences of the United States of America, 118(8). 10.1073/PNAS.1921481118

7. Calabrese, V., Santoro, A., Monti, D., Crupi, R., Di Paola, R., Latteri, S., Cuzzocrea, S., Zappia, M., Giordano, J., Calabrese, E. J., & Franceschi, C. (2018). Aging and Parkinson’s Disease: Inflammaging, neuroinflammation and biological remodeling as key factors in pathogenesis. In Free Radical Biology and Medicine (Vol. 115). 10.1016/j.freeradbiomed.2017.10.379

8. Di Benedetto, S., Müller, L., Wenger, E., Düzel, S., & Pawelec, G. (2017). Contribution of neuroinflammation and immunity to brain aging and the mitigating effects of physical and cognitive interventions. Neuroscience & Biobehavioral Reviews, 75, 114–128. 10.1016/j.neubiorev.2017.01.044

9. Franceschi, C., Bonafè, M., Valensin, S., Olivieri, F., De Luca, M., Ottaviani, E., & De Benedictis, G. (2000). Inflamm-aging. An evolutionary perspective on immunosenescence. Annals of the New York Academy of Sciences, 908(1), 244–254. 10.1111/j.1749-6632.2000.tb06651.x

10. Franceschi, C., Capri, M., Monti, D., Giunta, S., Olivieri, F., Sevini, F., Panourgia, M. P., Invidia, L., Celani, L., Scurti, M., Cevenini, E., Castellani, G. C., & Salvioli, S. (2007). Inflammaging and anti-inflammaging: A systemic perspective on aging and longevity emerged from studies in humans. Mechanisms of Ageing and Development, 128(1), 92–105. 10.1016/J.MAD.2006.11.016

11. Gilmer, G., Hettinger, Z. R., Tuakli-Wosornu, Y., Skidmore, E., Silver, J. K., Thurston, R. C., Lowe, D. A., & Ambrosio, F. (2023). Female aging: when translational models don’t translate. Nature Aging 2023 3:12, 3(12), 1500–1508. 10.1038/s43587-023-00509-8

12. Gitzlaff, L. A., Moody, J. F., Ma, Y., Johnson, S. C., Asthana, S., Betthauser, T. J., Christian, B. T., Alexander, A. L., Bendlin, B. B., Lauren Gitzlaff, C. A., & Alzheimer, W. (2024). Relationships between brain microstructure and Braak Stage progression on the Alzheimer’s disease continuum. Alzheimer’s & Dementia, 20, e095488. 10.1002/ALZ.095488

13. Guo, J., Huang, X., Dou, L., Yan, M., Shen, T., Tang, W., & Li, J. (2022). Aging and aging-related diseases: from molecular mechanisms to interventions and treatments. Signal Transduction and Targeted Therapy 2022 7:1, 7(1), 391-. 10.1038/s41392-022-01251-0

14. Hefendehl, J. K., Neher, J. J., Sühs, R. B., Kohsaka, S., Skodras, A., & Jucker, M. (2014). Homeostatic and injury-induced microglia behavior in the aging brain. Aging Cell, 13(1), 60–69. 10.1111/acel.12149

15. Herrera, M. L., Basmadjian, O. M., Falomir-Lockhart, E., Dolcetti, F. J. C., Hereñú, C. B., & Bellini, M. J. (2020). Sex frailty differences in ageing mice: Neuropathologies and therapeutic projections. European Journal of Neuroscience. 10.1111/ejn.14703

16. Horovitz, D. J., Askins, L. A., Regnier, G. M., & McQuail, J. A. (2025). Age-related synaptic signatures of brain and cognitive reserve in the rat hippocampus and parahippocampal regions. Neurobiology of Aging, 148, 80–97. 10.1016/j.neurobiolaging.2025.01.010

17. Ince, L. M., Darling, J. S., Sanchez, K., Bell, K. S., Melbourne, J. K., Davis, L. K., Nixon, K., Gaudet, A. D., & Fonken, L. K. (2023). Sex differences in microglia function in aged rats underlie vulnerability to cognitive decline. Brain, Behavior, and Immunity, 114, 438–452. 10.1016/J.BBI.2023.09.009

18. Knufinke, M., MacArthur, M. R., Ewald, C. Y., & Mitchell, S. J. (2023). Sex differences in pharmacological interventions and their effects on lifespan and healthspan outcomes: a systematic review. Frontiers in Aging, 4, 1172789. 10.3389/FRAGI.2023.1172789/FULL

19. Lemaître, J. F., Ronget, V., Tidière, M., Allainé, D., Berger, V., Cohas, A., Colchero, F., Conde, D. A., Garratt, M., Liker, A., Marais, G. A. B., Scheuerlein, A., Székely, T., & Gaillard, J. M. (2020). Sex differences in adult lifespan and aging rates of mortality across wild mammals. Proceedings of the National Academy of Sciences, 117(15), 8546–8553. 10.1073/PNAS.1911999117

20. López-Otín, C., Blasco, M. A., Partridge, L., Serrano, M., & Kroemer, G. (2023). Hallmarks of aging: An expanding universe. Cell, 186(2), 243–278. 10.1016/j.cell.2022.11.001

21. Mauvais-Jarvis, F., Arnold, A. P., & Reue, K. (2017). A Guide for the Design of Pre-clinical Studies on Sex Differences in Metabolism. Cell Metabolism, 25(6), 1216–1230. 10.1016/J.CMET.2017.04.033

22. Moldakozhayev, A., & Gladyshev, V. N. (2023). Metabolism, Homeostasis and Aging. Trends in Endocrinology and Metabolism: TEM, 34(3), 158. 10.1016/J.TEM.2023.01.003

23. Mosher, K. I., & Wyss-Coray, T. (2014). Microglial dysfunction in brain aging and Alzheimer’s disease. Biochemical Pharmacology, 88(4), 594–604. 10.1016/j.bcp.2014.01.008

24. Mota, C., Taipa, R., das Neves, S. P., Monteiro-Martins, S., Monteiro, S., Palha, J. A., Sousa, N., Sousa, J. C., & Cerqueira, J. J. (2019). Structural and molecular correlates of cognitive aging in the rat. Scientific Reports 2019 9:1, 9(1), 2005-. 10.1038/s41598-019-39645-w

25. Mottahedin, A., Ardalan, M., Chumak, T., Riebe, I., Ek, J., & Mallard, C. (2017). Effect of Neuroinflammation on Synaptic Organization and Function in the Developing Brain: Implications for Neurodevelopmental and Neurodegenerative Disorders. Frontiers in Cellular Neuroscience, 11(July), 1–16. 10.3389/fncel.2017.00190

26. Rattan, S. I. S. (2008). Hormesis in aging. Ageing Research Reviews, 7(1), 63–78. 10.1016/J.ARR.2007.03.002

27. Santoro, A., Martucci, M., Conte, M., Capri, M., Franceschi, C., & Salvioli, S. (2020). Inflammaging, hormesis and the rationale for anti-aging strategies. Ageing Research Reviews, 64, 101142. 10.1016/J.ARR.2020.101142

28. Sengupta, P. (2013). The Laboratory Rat: Relating Its Age with Human’s. In International Journal of Preventive Medicine (Vol. 4, Issue 6). www.ijpm.ir

29. von Bernhardi, R., Eugenín-von Bernhardi, L., & Eugenín, J. (2015). Microglial cell dysregulation in brain aging and neurodegeneration. Frontiers in Aging Neuroscience, 7(JUN), 1–21. 10.3389/fnagi.2015.00124

